# Spontaneous brain activity and synaptic density in schizophrenia: a combined [11C]UCB-J PET and fMRI study

**DOI:** 10.1101/2024.09.25.614893

**Authors:** Ekaterina Shatalina, Ellis Chika Onwordi, Thomas Whitehurst, Alexander Whittington, Ayla Mansur, Atheeshaan Arumuham, Tiago Reis Marques, Roger N. Gunn, Sridhar Natesan, Eugenii A. Rabiner, Matthew B. Wall, Oliver D Howes

## Abstract

Schizophrenia is associated with altered Amplitude of Low Frequency Fluctuations (ALFF), a functional Magnetic Resonance Imaging (fMRI) measure of spontaneous brain activity at rest. ALFF in healthy controls has been linked with presynaptic density levels measured by [11C]UCB-J positron emission tomography (PET). Given the growing body of evidence for low presynaptic density levels in schizophrenia, we set out to test if altered [11C]UCB-J binding may be associated with changes in ALFF in schizophrenia, and secondly to test whether the relationships between ALFF and [11C]UCB-J binding differ at the group level.

Subjects with schizophrenia had higher ALFF in the medial prefrontal cortex and other regions, in line with published meta-analyses. In control subjects, there was a significant positive relationship between [11C]UCB-J distribution volume ratio (DVRcs) and ALFF in the medial prefrontal cortex (r=0.54, p=0.0365, n=16), but not in subjects with schizophrenia (r=–0.14, p=0.5564, n=22); r-coefficients significantly differed between groups (Z_observed_=2.07, p=0.019). At the whole brain level, there were significant positive correlations between [11C]UCB-J DVRcs and ALFF in control subjects in the putamen, insular cortex, precentral gyrus and occipital regions, while in the schizophrenia group, there were significant positive correlations in the bilateral dorsolateral prefrontal cortex and negative correlations in the cuneus, parietal lobule and supramarginal gurus. Correlation coefficients were significantly different between groups across all cortical and subcortical regions with both higher and lower correlation coefficients in the control group.

Our results suggest a link between spontaneous brain activity and presynaptic density in control subjects and that this relationship may be disrupted in schizophrenia patients, despite higher ALFF in this group, indicating altered neurobiological mechanisms. Widespread significant differences in ALFF-[11C]UCB-J DVRcs correlation coefficients between controls and schizophrenia subjects highlight the complexity of synaptic dysfunction in schizophrenia and underscore the need for further research to explore the underlying biological mechanisms.

## Introduction

Schizophrenia is a common and disabling neuropsychiatric disorder associated with positive (e.g. psychosis, delusions, hallucinations), negative (e.g. flattening of emotions, anhedonia) and cognitive symptoms (impaired executive function, impaired social cognition). A better understanding of the brain alterations underlying schizophrenia is critical for developing improved treatments [1].

Schizophrenia is characterised by several alterations in brain function, including altered Amplitude of Low Frequency Fluctuations [2]. ALFF is a measure of brain function derived from the Blood Oxygen Level Dependent signal measured during functional Magnetic Resonance Imaging (fMRI). It is typically defined as the power of the fluctuations in the BOLD signal in the 0.01-0.1Hz frequency range and is thought to reflect spontaneous neural activity during a resting state scan [3, 4]. Upon meta-analysis, patients with schizophrenia have altered ALFF and fractional ALFF (fALFF), which is ALFF normalised to the entire frequency range of the signal time series, when compared to matched controls. Specifically, schizophrenia is associated with significantly lower (f)ALFF in the somatosensory cortex, posterior parietal cortex, and occipital cortex and higher (f)ALFF in the bilateral striatum, medial temporal cortex, and medial prefrontal cortex relative to controls [2].

Some recent work has shown that, in healthy adults, presynaptic terminal density levels measured using [11C]UCB-J positron emission tomography (PET) are related to resting state fALFF. Specifically, greater medial prefrontal presynaptic terminal density was associated with greater fALFF of the anterior default mode, posterior default mode, and executive control networks, while greater striatal synaptic density was associated with greater fALFF of the anterior default mode and salience networks [5]. *In vivo* imaging data suggest individuals with chronic schizophrenia have lower levels of a presynaptic terminal marker in the frontal cortex and the anterior cingulate cortex, as well as in other brain regions, and with evidence that differences may be both more or less pronounced earlier in illness [6–9]. These studies raise the question of whether lower presynaptic terminal density may underlie differences in brain function associated with schizophrenia. Given these findings, and that BOLD contrast is thought to reflect synaptic activity [10–17], testing whether differences in presynaptic terminal density associated with schizophrenia are associated with changes in brain function is a key question to address.

We tested whether differences in [11C]UCB-J binding, which can be interpreted as a measure of presynaptic terminal density, may explain the aberrant ALFF reported in schizophrenia. Studies in schizophrenia have used both ALFF and fALFF. However, while fALFF is a useful method of reducing noise in ALFF signal, it may not be suitable for comparing groups where the frequency range could be altered by a disease condition or drug and it has been shown to be less reliable during test-retest validation [18]. We therefore investigated the relationship between ALFF and SV2A levels in unmedicated patients with schizophrenia and healthy control subjects, to test whether differences in presynaptic terminal levels may underlie aberrant ALFF in patients and secondly to test whether the relationships between ALFF and [11C]UCB-J binding differ at the group level.

## Methods

### 2.1 Approvals and recruitment

The study was approved by the West London & GTAC Research Ethics Committee (16/LO/1941) and was conducted in compliance with the principles of Good Clinical Practice (GCP), the Declaration of Helsinki (1996 Version), the Research Governance Framework for Health & Social Care, and the Administration of Radioactive Substances Advisory Committee (ARSAC) guidelines. Standard MRI screening procedures were followed and written informed consent was obtained from all participants.

Inclusion criteria for all subjects were: aged 18 to 65 years old, having adequate command of English, and a normal blood coagulation test. Exclusion criteria were: drug or alcohol dependence (except nicotine), physical illness, past or present neurological illness (excluding schizophrenia for the patient group), pregnancy or lactating mothers, imaging contraindications and taking a drug known to interact with SV2A (including levetiracetam, brivaracetam, loratadine or quinine [19]. For healthy volunteers, additional exclusion criteria were current or past psychiatric diagnosis or family history of schizophrenia.

Control subjects were recruited through advertising and patients were recruited from London first-episode services. Patient-specific inclusion criteria were meeting DSM-5 criteria for schizophrenia and being antipsychotic-naïve or free from any antipsychotic medication for at least 4 weeks before imaging. Clinical assessments included the Structured Clinical Interview for DSM-5 and the Positive And Negative Symptoms Scale (PANSS) [20]. Illness duration was determined as the time from each patient’s first psychotic symptoms.

### 2.2 Magnetic resonance imaging acquisition

Data were acquired on a Siemens MAGNETOM Prisma 3 Tesla (3T) MRI scanner (Siemens Healthineers, Erlangen, Germany) with the in-built body coil used for Radio Frequency (RF) excitation and the manufacturer’s 64-channel phased-array head/neck coil for reception.

Whole-head anatomical T1-weighted images were acquired at the beginning of each scanning session using the Magnetisation Prepared Rapid Gradient Echo (MPRAGE) sequence, with parameters based on Alzheimer’s Disease Research Network (ADNI-GO); FOV 256 x 256mm, 1mm isotropic voxels, 176 sagittal slices, repetition time (TR) = 2300ms, echo time (TE) = 2.98ms, Inversion time = 900ms, flip angle 9°, bandwidth 200Hz/pixel, Parallel Imaging factor (PI) of 2, [21]. All structural images were inspected by an experienced clinical neuroradiologist for unexpected findings of clinical significance. If such findings were identified, participants were excluded from the study.

Functional data were acquired for a duration of 8 minutes and consisted of T2* weighted transverse echo planar image (EPI) slices. A total of 190 volumes was collected in an ascending direction with 3.4×3.4×3.0mm voxels, FoV = 220mm, TR= 2500 ms, TE = 30ms, total slices = 42, Flip angle = 90°, Bandwidth 1594 Hz/Px, GRAPPA = 2. Participants were given instructions to lay still with their eyes open and not think about any one thing and let their minds wander.

### 2.3 Data pre-processing

All functional data and anatomical data were processed with FSL (FMRIB Software Library v5.0.4; http://www.fmrib.ox.ac.uk/fsl/). BET was used for brain extraction of the anatomical data and the fsl_anat script was used for additional anatomical data pre-processing. Motion correction was performed with FMRIB Linear Image Registration Tool (MCFLIRT), with spatial smoothing using a Gaussian kernel of full width at half maximum (FWHM) 6mm. A two-step co-registration was performed, first to the subject’s individual anatomical image followed by registration to an anatomical template image in standard stereotactic space (MNI152), with no temporal filtering applied at this stage to ensure the data captured the full frequency range. Mean white matter and CSF signals were used as regressors of no interest alongside six head motion regressors. High motion for a subject was defined as a mean relative root-mean square displacement that exceeds 0.5 mm; no subjects included in this study exceeded this bound.

### 2.4 Amplitude of low frequency fluctuations analysis

Amplitude of low frequency fluctuations (ALFF) was calculated using the Analysis of Functional Neuroimages (AFNI v.20.1.06) 3dRSFC module. The data in these analyses were band-pass filtered using a range of 0.01-0.1Hz and results were converted to Z-statistics for second-level analyses. As ALFF is sensitive to the raw values of the BOLD time series (which are essentially arbitrary values) some normalisation of ALFF measures is standard practice [22]. In this case, *Z*-normalisation was used where the mean of each voxel’s time series was divided by its standard deviation, to give a *Z*-score for that voxel, and an overall Z-score brain map for each participant.

### 2.5 Second-level analyses

Second-level analyses were carried out at the whole-brain level. ALFF images were combined using FSL’s FLAME-1 model to compare patients with schizophrenia and control subjects. Results were thresholded at Z=2.3 and a cluster threshold of p<0.05 (corrected for multiple comparisons). A spherical region of interest (ROI) of a 10mm radius was formed around the peak of the ALFF second-level analyses capturing the region where differences in ALFF were greatest between groups. This ROI was used to extract both ALFF and [11C]UCB-J DVR_CS_ values for each subject. Partial correlation was used to assess relationships between ALFF and [11C]UCB-J DVR_CS_ in each group in this region, with age included as a control variable, as schizophrenia and control groups were significantly different in age (table 1). Resultant r-coefficients were compared using the Z-observed method detailed in section 2.6 below. Outlier detection was implemented by transforming data for each group to Z-scores, and data points with a Z-score exceeding ±3 were classified as outliers [23]. An independent t-test was used to compare [11C]UCB-J DVR_CS_ values between the two groups in this region.

**Table 1:**
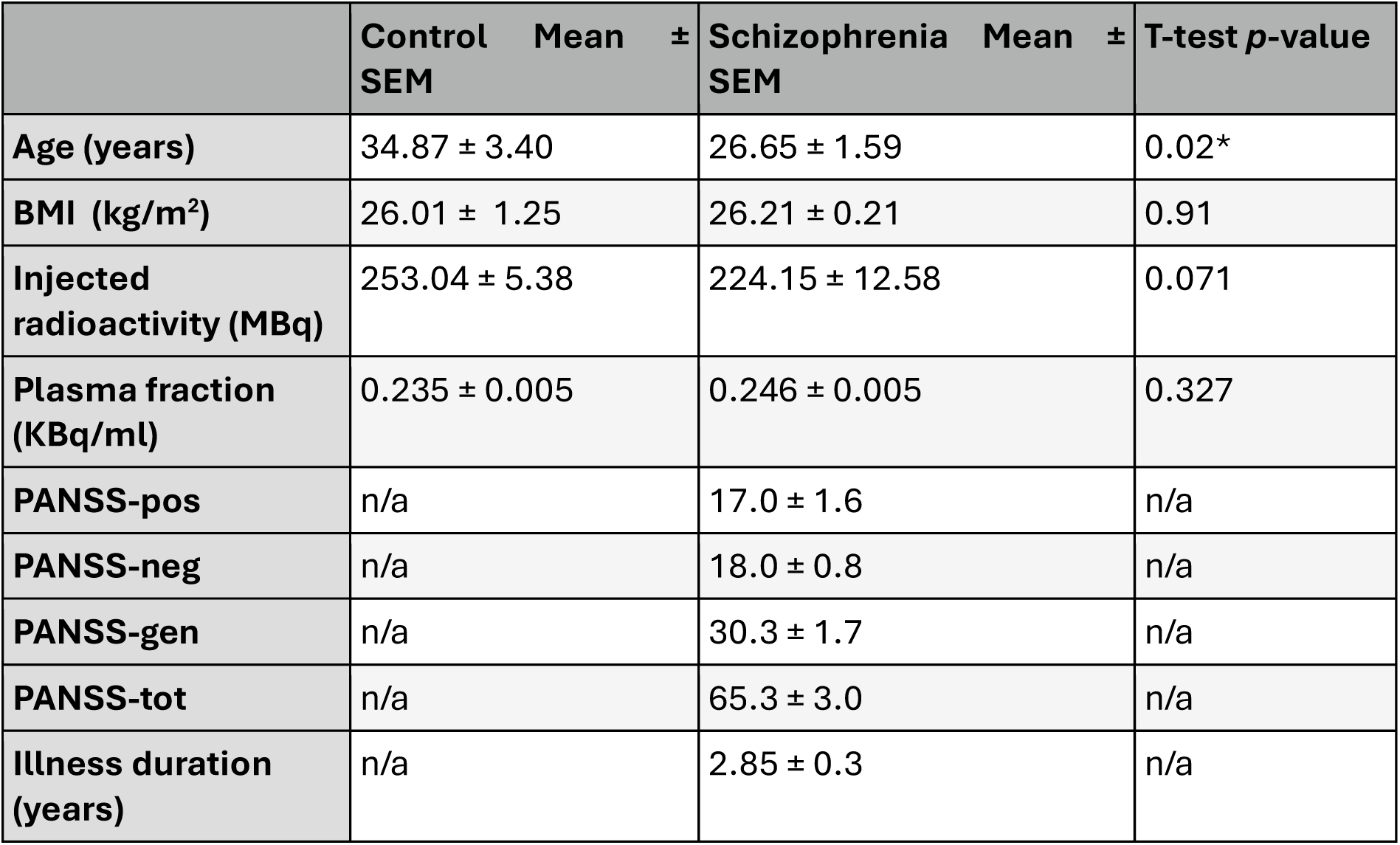
Demographic details

### 2.6 [11C]UCB-J data acquisition and processing

[11C]UCB-J data was acquired as described previously by Onwordi et al., (2024) on a HiRez 6 PET/computed tomography scanner (Siemens Healthcare, Erlangen, Germany) [7]. The tracer was administered as an intravenous bolus injection over 20s into the cubital vein, followed by data being collected continuously for 90 min over 26 frames (frame durations: 8 × 15 s, 3 × 60 s, 5 × 120 s, 5 × 300 s and 5 × 600 s). A unilaterally placed arterial line was used to collect arterial blood samples. From these the arterial input function was calculated, which included the associated processing of discrete samples to determine the plasma-to-whole blood ratio and the parent fraction.

MIAKAT (https://nmmitools.org/2019/01/01/miakat/) was used for processing and modelling the data. MIAKAT is implemented using MATLAB (version R2017a; The MathWorks, Inc., Natick, Mass.) and uses FSL (version 4.1.9) functions for brain extraction, and SPM12 (http://www.fil.ion.ucl.ac.uk/spm) functions for image segmentation and registration.

Each PET frame was adjusted for radioactive decay and head movement using rigid-body registration, with the 16th frame serving as the reference. Additionally, the T1-weighted MR image underwent co-registration with the summed PET image, following the extraction of the brain using the Brain Extraction Tool. [24].

Analysis was carried out using the 1-tissue compartment (1TC) tissue model using metabolite-corrected plasma input fraction with a fixed 5% blood volume correction, as this has been previously validated for this tracer [25, 26]. All voxel time-activity curves were analyzed in subject space to derive a distribution volume (V_T_; mL/cm^3^) for each voxel, similar to previous work [27, 28]. Non-linear deformation parameters were derived by the diffeomorphic anatomical registration through exponentiated lie algebra (DARTEL) algorithm and used for the mapping of the T1 image into stereotaxic space, to transform each subject’s PET data into standard MNI152 space following kinetic modelling [29].

The distribution volume ratio (DVR_CS_) was used as the main outcome measure. To calculate DVRcs, a mask of the centrum semiovale (CS) region was used to extract an average CS V_T_ value for each subject. Following this, each subject’s parametric map was divided by their CS V_T_, to produce a parametric DVR_CS_ map. DVR_CS_ has been validated as an optimal outcome measure for this tracer in previous work [26, 28].

### 2.7 Exploratory whole-brain UCB-J PET and resting state fMRI analyses

We conducted whole-brain correlation analyses in control subjects and in patients with schizophrenia to explore if there were relationships in other regions, as previously implemented in [30]. Separately for each group, parametric ALFF Z-statistic images and respective [11C]UCB-J DVR_CS_ images were concatenated into 4d datasets using FSL’s fslmerge tool and masked using a 75% grey matter mask derived from the MNI152 T1 structural template. AFNI’s 3dTcorrelate module was then used to run voxel-wise Pearson’s correlations between the PET and fMRI datasets, correlating each voxel’s SV2A distribution values (across subjects) with the ALFF Z-statistics.

Resultant *r*-coefficient images were transformed to *t*-statistical images using fslmaths, using formula 1 below, and these t-maps were then transformed to *Z*-scores using FSL’s ‘ttoz’ function. FSL’s ‘easythresh’ function was then used to threshold the images at *Z*=2.3, with cluster extent thresholding applied at *p*<0.05 to correct for multiple comparisons across the brain. A threshold of *Z*=2.3 was used as these analyses are of an exploratory nature. Final figures were made by creating a binary mask of significant clusters for each analysis and using this to mask the original *r*-images to show the *r*-coefficients corresponding to the significant findings. Unthresholded images are provided in the supplement as an exploratory measure of the interrelationship between [11C]UCB-J DVR_CS_ and ALFF. To compare r-coefficients between groups in a voxel-wise manner, *Z*-scores for each group were used to calculate Z_observed_ values using formula 2 and fslmaths, following which FSL’s ‘easythresh’ function was then used to threshold the images at *Z*=3.1 with cluster extent thresholding applied at *p*<0.05 to correct for multiple comparisons across the brain.

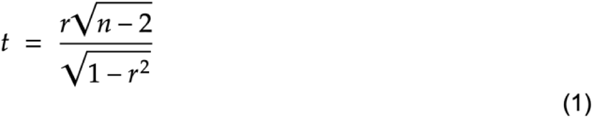

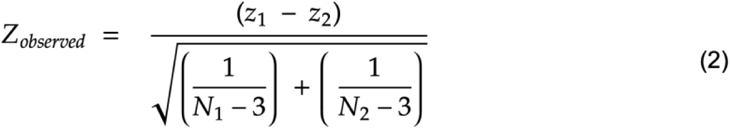

## Results

### Group differences in ALFF and relationship with [11C]UCB-J DVRcs

A whole-brain comparison of subjects with schizophrenia and controls showed significantly higher ALFF values in the patient group across several regions (FLAME-1, Z>2.3, cluster p<0.05, figure 1). These differences were localised to frontal cortical regions, the anterior cingulate and the caudate, asshown in figure 1. There were no areas of the brain where ALFF was significantly lower in patients at the whole brain level. The peak in group differences was observed in the medial frontal cortex (*Z*=3.88, MNI x=44, y=86, z=29) and was used to form a spherical ROI (figure 2A). There was no significant difference in [11C]UCB-J DVR_CS_ between the groups in this region (t(38)=1.757, *p*=0.087, two-sided independent-test), shown in Figure 2B, although in absoluteterms it waslower in the schizophrenia group.

**Figure 1:**
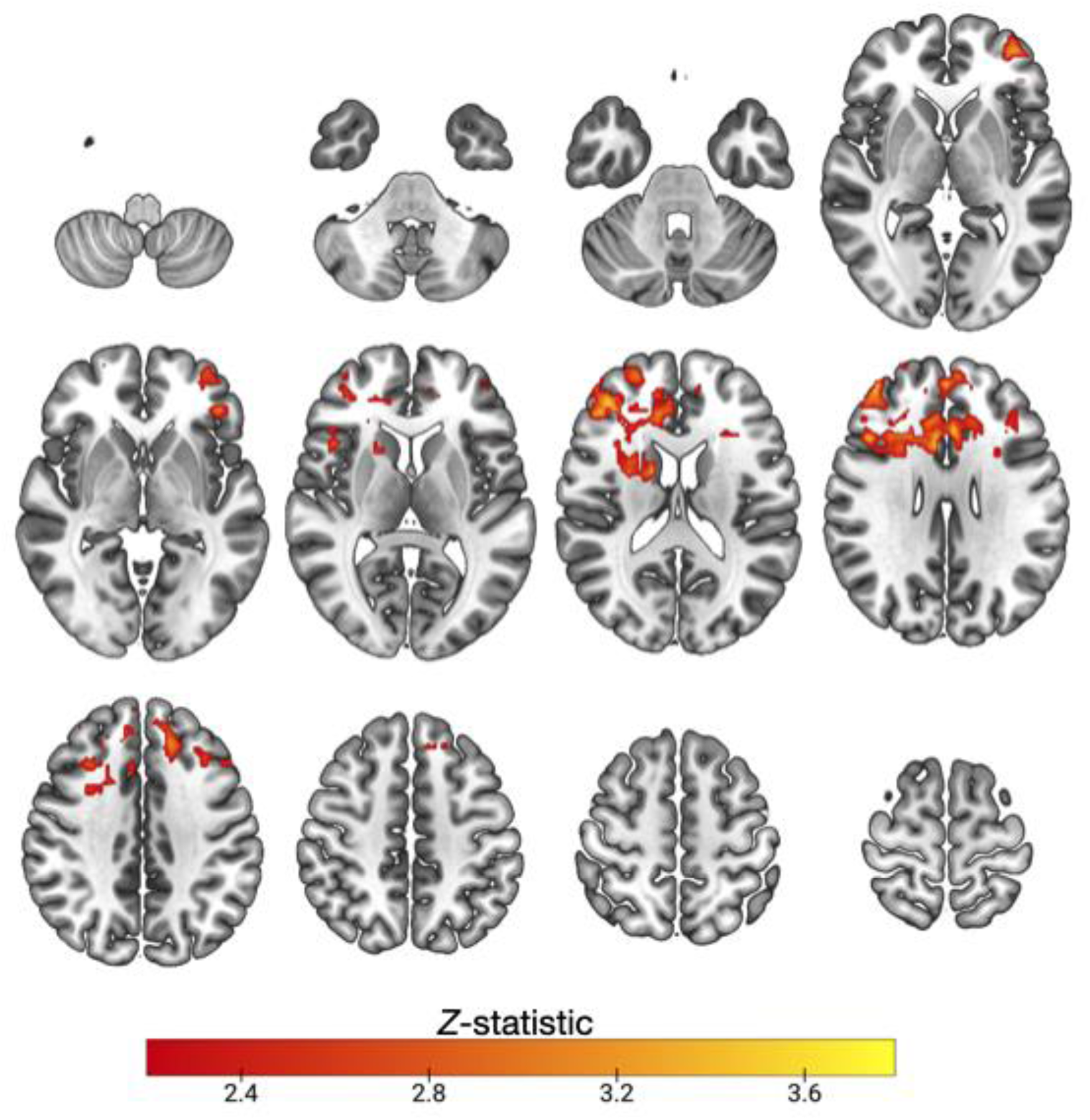
Significantly higher resting state amplitude of low frequency fluctuations (ALFF) in unmedicated subjects with schizophrenia: Z (Gaussianised T/F) statistical images representative of significant differences in amplitude of low frequency fluctuations at rest in unmedicated patients with schizophrenia (*n*=22) compared to healthy controls (*n*=16). Images were thresholded using clusters determined by *Z* > 2.3 and a (corrected) cluster significance threshold of *p* = 0.05. Axial slices shown in MNI152 are: -52 -42 -32 -22-12; -2 8 18 28; 38 48 56 66, images are in neurological orientation L=L.

**Figure 2:**
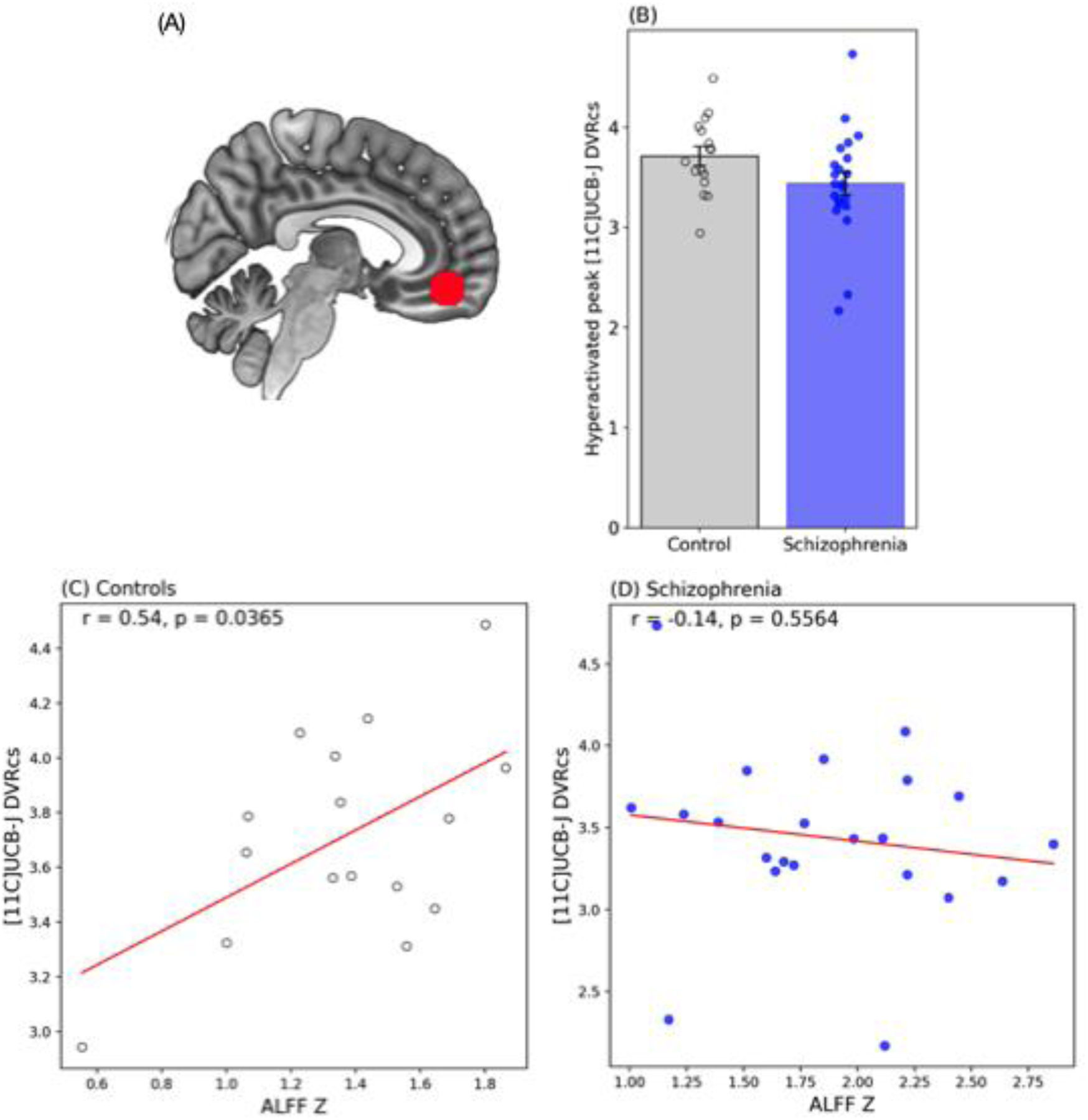
Higher amplitude of low frequency fluctuations (ALFF) in unmedicated subjects with schizophrenia and corresponding UCB-J DVR values: (A) Mask showing region of interest (ROI) formed around the peak group difference in ALFF, where schizophrenia subjects had higher ALFF than control subjects. MNI152 x=-35, y=17, z=27. (B) Mean ROI [11C]UCB-J DVR_CS_ in patients with schizophrenia and respective controls (independent t-test, p=0.087). (C) Scatter plot showing the relationship between [11C]UCB-J DVR_CS_ and ALFF Z-statistics in the hyperactivated cluster, showing a significant positive relationship in controls but not people with schizophrenia (D). Trend lines mark partial correlation results for control subjects (hollow black data points, *n* =16) and for patients with schizophrenia (blue data points, *n* = 22).

Figure 2C shows that control subjects had a significant positive correlation between [11C]UCB-J DVRcs and ALFF in the medial frontal cortex (Partial correlation, r=0.54, p=0.0365), while subjects with schizophrenia did not have a significant relationship between the two measures in this region (Partial correlation, r=–0.14, p=0.5564). There were no outliers in any group. Given that the groups were significantly different in age (table 1), age was included as a covariate in the partial correlation. The r-coefficients were significantly different between the two groups (Z_observed_=2.07, p=0.019). There were no significant associations between ALFF and PANSS scores or between ALFF and illness duration in the schizophrenia group (supplementary table 1).

### Exploratory whole-brain analysis of relationships between ALFF and [11C]UCB-J DVRcs

Whole brain correlation analyses showed that in control subjects [11C]UCB-J DVR_CS_ was significantly positively correlated with ALFF in regions spanning the putamen, insular cortex and precentral gyrus, angular gyrus, cerebellum, visual areas, and dorsal occipital regions; see figure 3A. Patients with schizophrenia showed a different pattern of associations between [11C]UCB-J DVRcs and ALFF which included significant positive associations in the bilateral dorsolateral prefrontal cortex and brainstem, with negative associations in the cuneus, parietal lobule and supramarginal gyrus (figure 3B). Unthresholded correlation maps for the control and schizophrenia groups are provided in the supplement for reference (supplementary figures 1 and 2, respectively). An analysis comparing *r*-coefficients between control subjects and patients with schizophrenia showed that control subjects had significantly higher r-coefficients than the schizophrenia group in regions which included parts of the occipital, parietal, and frontal medial cortices (shown in red, figure 4). The control group had significantly lower r-coefficients than the schizophrenia group in regions including parts of the cerebellum, subcortical areas including the putamen and thalamus and DLPFC (shown in blue, figure 4).

**Figure 3:**
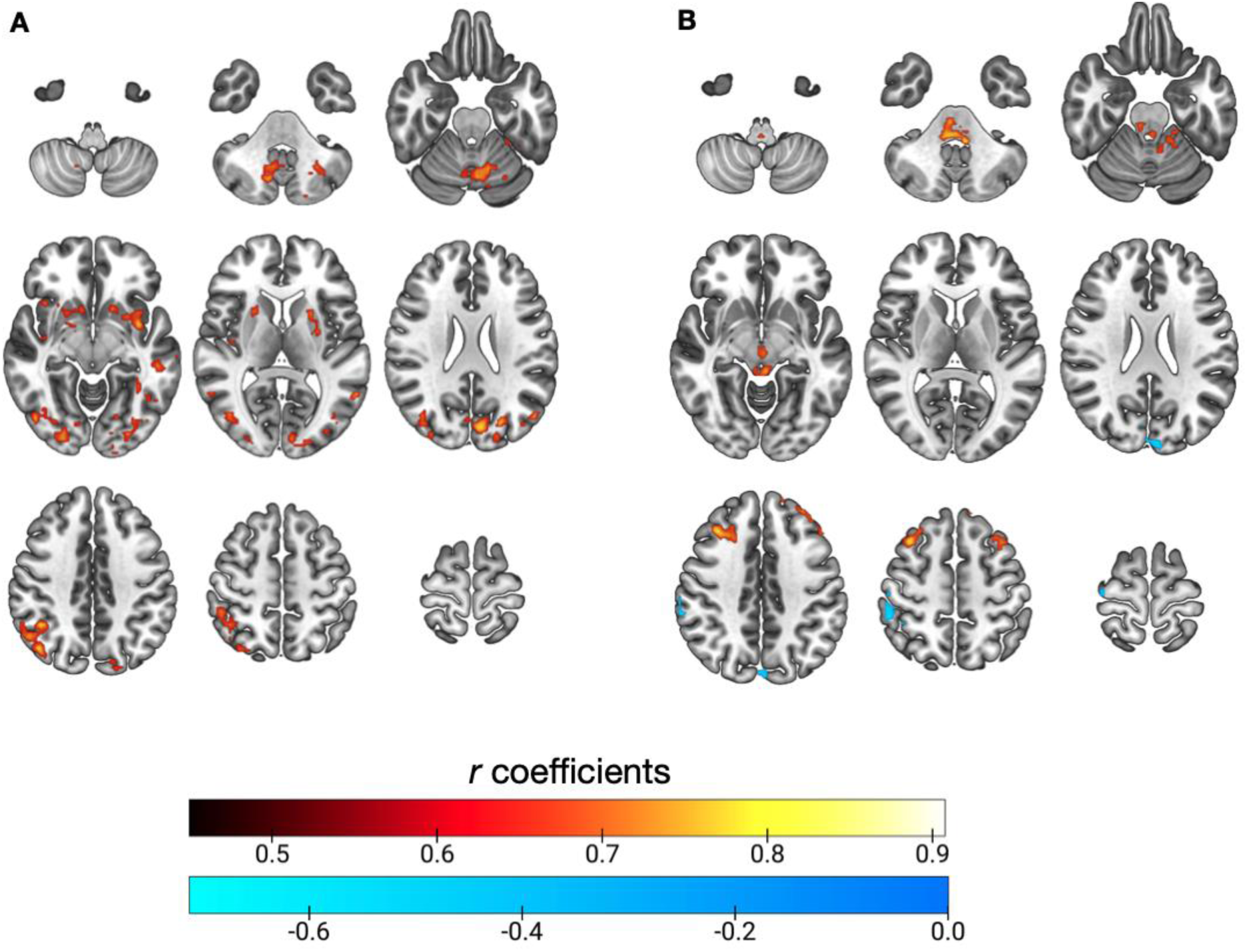
Significant brain-wide correlations between resting state amplitude of low frequency fluctuations and [11C]UCB-J DVRcs in (A) control subjects and (B) subjects with schizophrenia : r-coefficient statistical images representative of significant brain-wide correlations between amplitude of low frequency fluctuations at rest and [11C]UCB-J DVR_CS_. Healthy control subjects *n*=16, unmedicated subjects with schizophrenia (*n*=22). Voxel-wise Pearson’s correlations, transformed to Z-statistical images thresholded at *Z*=2.3 and a cluster-wise threshold of *p*<0.05. Axial slices shown in MNI152 are: A -50 -40 -24; -8 8 24; 40 54 70, images are in neurological orientation L=L.

**Figure 4:**
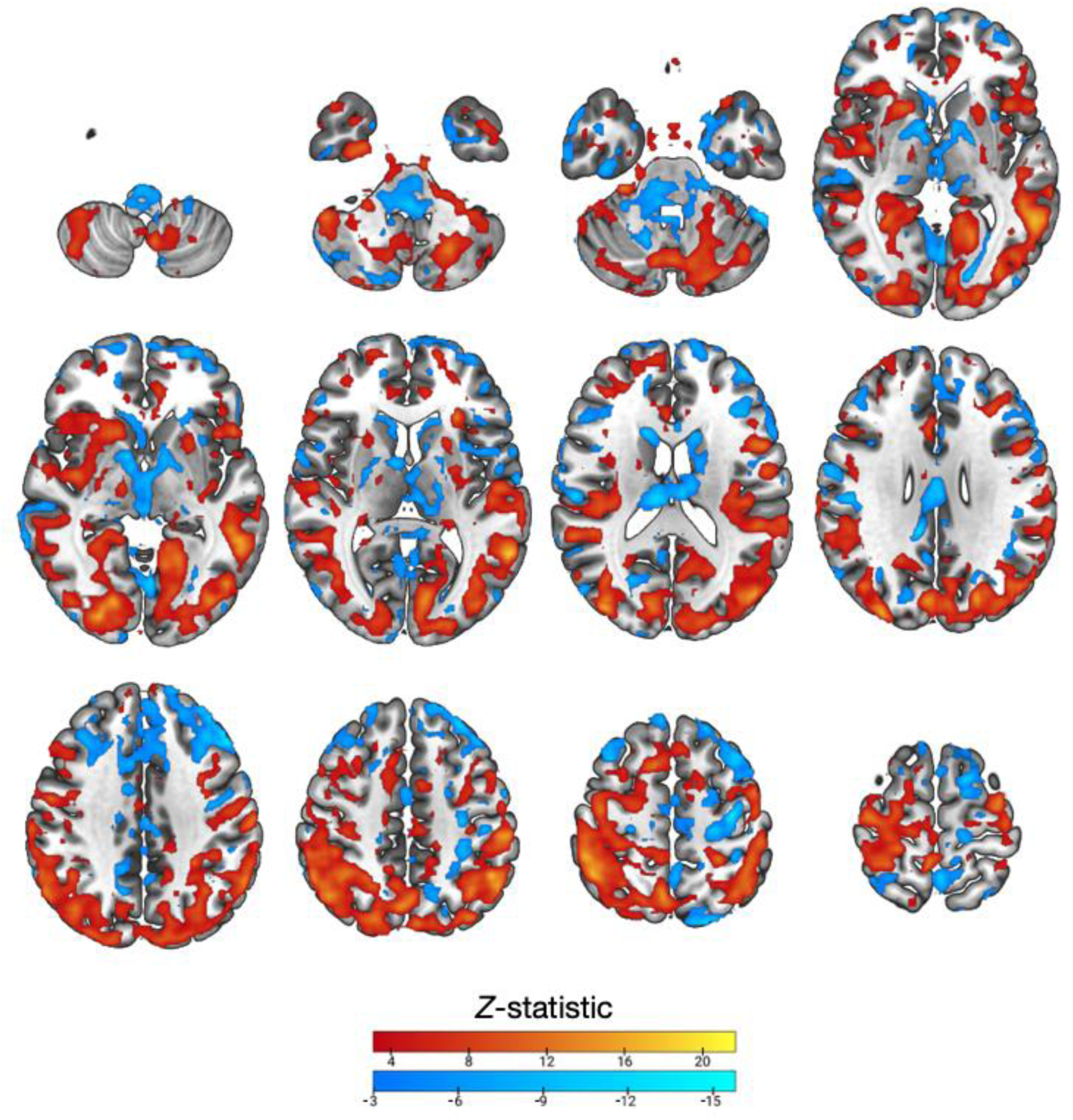
Significant differences in brain-wide correlations between resting state amplitude of low frequency fluctuations and [11C]UCB-J DVRcs between control subjects and patients with schizophrenia: Z-statistical images thresholded at *Z*=3.1 and a cluster-wise threshold of *p*<0.05 comparing r-coefficients for associations between amplitude of low frequency fluctuations at rest and [11C]UCB-J DVR_CS_ in healthy control subjects (*n*=16) and patients with schizophrenia (*n*=22). Blue shows significantly lower *r*-values in controls, and yellow-red shows significantly higher *r*-values in controls. Axial slices shown in MNI152 are: -52 -42 -32 - 22-12; -2 8 18 28; 38 48 56 66, images are in neurological orientation L=L.

## Discussion

The main finding of this study is that in subjects with schizophrenia ALFF in the medial prefrontal cortex was not significantly related to [11C]UCB-J DVRcs, and the correlation coefficient in the patient group was significantly lower than the control group. In contrast, in healthy controls, ALFF was significantly positively related to [11C]UCB-J DVRcs in this same region. Results in healthy controls are in line with previous work by Fang et al., with significant positive correlations between [11C]UCB-J volume of distribution (*V*_T_) in the medial prefrontal cortex and fALFF of the anterior default mode, posterior default mode, and executive control networks [5]. Given that in this region patients with schizophrenia had significantly higher ALFF, but comparable [11C]UCB-J DVRcs, these findings suggest the ALFF–[11C]UCB-J DVRcs relationship may be disrupted in schizophrenia, but not in a manner that was linearly related to an aberrant increase in ALFF.

Our findings of higher ALFF in the medial frontal cortex, anterior cingulate and caudate in patients with schizophrenia compared with controls, are consistent with a recent meta-analysis which found increased ALFF in the inferior frontal gyrus and anterior cingulate in a large sample of 1249 schizophrenia patients and 1179 controls [2]. This is also partially in line with another recent meta-analysis of ALFF/fALFF in drug-naïve first-episode patients, which reports increased ALFF/fALFF in the putamen, posterior cingulate cortex, cerebellum, middle frontal gyrus and superior frontal gyrus [31]. Given that our study was carried out in unmedicated patients, antipsychotic medication could not have influenced the findings.

Our finding in control subjects that [11C]UCB-J DVR_CS_ was significantly positively correlated with ALFF in regions spanning the putamen, insular cortex and precentral gyrus, angular gyrus, cerebellum, visual areas and occipital regions is in agreement with previous work published by Fang et al. (2023) where the authors report correlations between striatal [11C]UCB-J binding with fALFF in the motor and salience networks [5, 32]. Interestingly, at the whole-brain level, ALFF–[11C]UCB-J DVRcs correlation coefficients were significantly and strikingly different between patients with schizophrenia and controls, across large portions of the brain. This included both areas where the strength of association between SV2A density and ALFF was higher as well as lower in patients with schizophrenia. Together these significant differences spanned most cortical and subcortical regions.

### Strengths and limitations

Before discussing biological interpretations, it is also important to consider and address methodological considerations, strengths and limitations relevant for interpreting these findings. One key strength of this study is the joint use of BOLD fMRI and [11C]UCB-J PET in the same subjects. Furthermore, one advantage of using ALFF over other approaches for evaluating brain function at rest, is that it does not depend on defining activity in the context of the activity of another brain region (i.e. functional connectivity), which is a key limitation of correlation-based or independent components analysis (ICA) methods [17]. However, while BOLD signal has been shown to reflect local field potentials, suggesting it captures synaptic firing [33], there is outstanding discussion around the biological interpretation of resting state BOLD signal and therefore of ALFF [34].

Another strength of this study is the inclusion of only unmedicated subjects with schizophrenia, which reduces the possible confounding effects of antipsychotics on measures of brain function or on [11C]UCB-J binding. It is worth noting that differences in presynaptic terminal density levels in this sample are less pronounced when compared to matched controls [7], which makes it especially interesting to test if our findings translate to chronic schizophrenia patients who have lower [11C]UCB-J binding and greater across-subject variability in [11C]UCB-J DVRcs [6].

There are also several key considerations. Firstly, ALFF is a measure that is sensitive to physiological noise and head motion, and cohorts with psychiatric diagnoses are known to have higher head motion than healthy controls [35]. Importantly, none of the subjects included in this study had high head motion, which makes it unlikely that motion would explain the brain-wide differences in ALFF–[11C]UCB-J DVRcs relationships, however, it is possible that subtle differences in motion could impact the ALFF measure. Additionally, the small sample size of this study is a key consideration, especially the limited size of the control group, as it constrains the conclusions, we can draw from the group differences. Given the small sample size, singular data points could also have a significant impact on the correlation strength and significance. While our main findings are broadly in line with previous work [5], Fang et al., (2024) used an overlapping, but not an identical ROI to the one we defined in this study. This highlights the importance of replicating our findings in an independent cohort.

### Interpretation of findings and implications

[11C]UCB-J PET specifically binds the SV2A protein, which is ubiquitously expressed on vesicles across all synapse types [36, 37]. Its levels have been shown to correlate with that of synaptophysin [37], which is considered the gold standard *in vitro* molecular marker of synaptic density [38, 39]. The field has been widely interpreting [11C]UCB-J PET as a marker of synaptic density [40], although a more precise interpretation would be a marker of presynaptic terminal density given the presynaptic origin of the protein.

ALFF is derived from the BOLD contrast time series and is the power of the Fourier-transformed time series in the 0.01-0.1 Hz frequency range. Studies employing the measure have interpreted ALFF as a measure of spontaneous brain activity at rest [4, 41]. Several studies have shown that (f)ALFF is positively spatially correlated with glucose utilisation measured using fludeoxyglucose (FDG) PET in healthy subjects and clinical groups [42–45]. Given that synaptic firing is highly energy intensive, these data suggest ALFF may be capturing spatial variation in synaptic metabolism at the subject level. However, studies relating the two measures across subjects have produced mixed results [34, 42, 43, 45] which warrants further investigation into whether ALFF quantitatively captures variation in neuronal metabolic activity. Similarly, several studies have shown correlations between ALFF and cerebral blood flow [46], while others have also shown there are regions where there is no spatial overlap between across-subject variation in the two measures [47]. While this evidence paints a somewhat mixed picture, the most recent work from Fang et al. (2024) showing robust across-subject correlations between fALFF and [11C]UCB-J VT, in a relatively large cohort, provides a compelling interpretation of ALFF as a functional measure with a possible synaptic origin.

Positive relationships between [11C]UCB-J DVRcs and ALFF seen in our control group are in line with ALFF having a synaptic origin. Disrupted relationships in the schizophrenia group, notably the presence of significant negative correlations between ALFF and [11C]UCB-J DVRcs, which are absent in the control group, may have several biological interpretations. For example, a large body of literature indicates schizophrenia is associated with excitation-inhibition imbalance [48], which could mean that the contribution of inhibitory and excitatory synapses to total synaptic density would differ between schizophrenia and controls, even in the context of comparable presynaptic density. In line with this, recent work combining [11C]UCB-J PET imaging with magnetic resonance spectroscopy suggests relationships between [11C]UCB-J DVR_CS_ and glutamate levels are disrupted in schizophrenia when compared to controls [49]. In our present study, we found greater presynaptic terminal density in the cuneus was associated with lower ALFF in schizophrenia subjects. This raises the question of whether differences in the ratios of inhibitory and excitatory synapses in this region in schizophrenia may contribute to these results. However, additional experiments would be required to test this hypothesis with careful consideration of the differences in metabolic demand between glutamatergic and GABAergic neurons [50] and their subsequent contributions to the haemodynamic response measured by the BOLD contrast.

### Conclusions

Our study showed that in subjects with schizophrenia, ALFF in the medial prefrontal cortex was not significantly related to [11C]UCB-J DVRcs, while in healthy controls there was a significant positive relationship, which indicates altered neurobiological mechanisms in schizophrenia. We found widespread significant differences in ALFF-[11C]UCB-J DVRcs correlations between controls and schizophrenia patients, spanning a wide range of cortical and subcortical regions. Together these findings highlight the complexity of synaptic dysfunction in schizophrenia and underscore the need for further research to explore the underlying biological mechanisms.

### Declarations

#### Data Availability statement

The datasets used and/or analysed during the current study are available from the corresponding author upon reasonable request.

## Author contributions

Study concept and design: ES, MW, OH

Data collection: ES, TW, ECO

Acquisition, analysis, or interpretation of data: all authors

Drafting of the manuscript: all authors

Critical revision of the manuscript for important intellectual content: all authors

Statistical analysis: ES

Study supervision: OH, MW

## Acknowledgements

Not applicable

## Supplementary material

**Supplementary figure 1:**
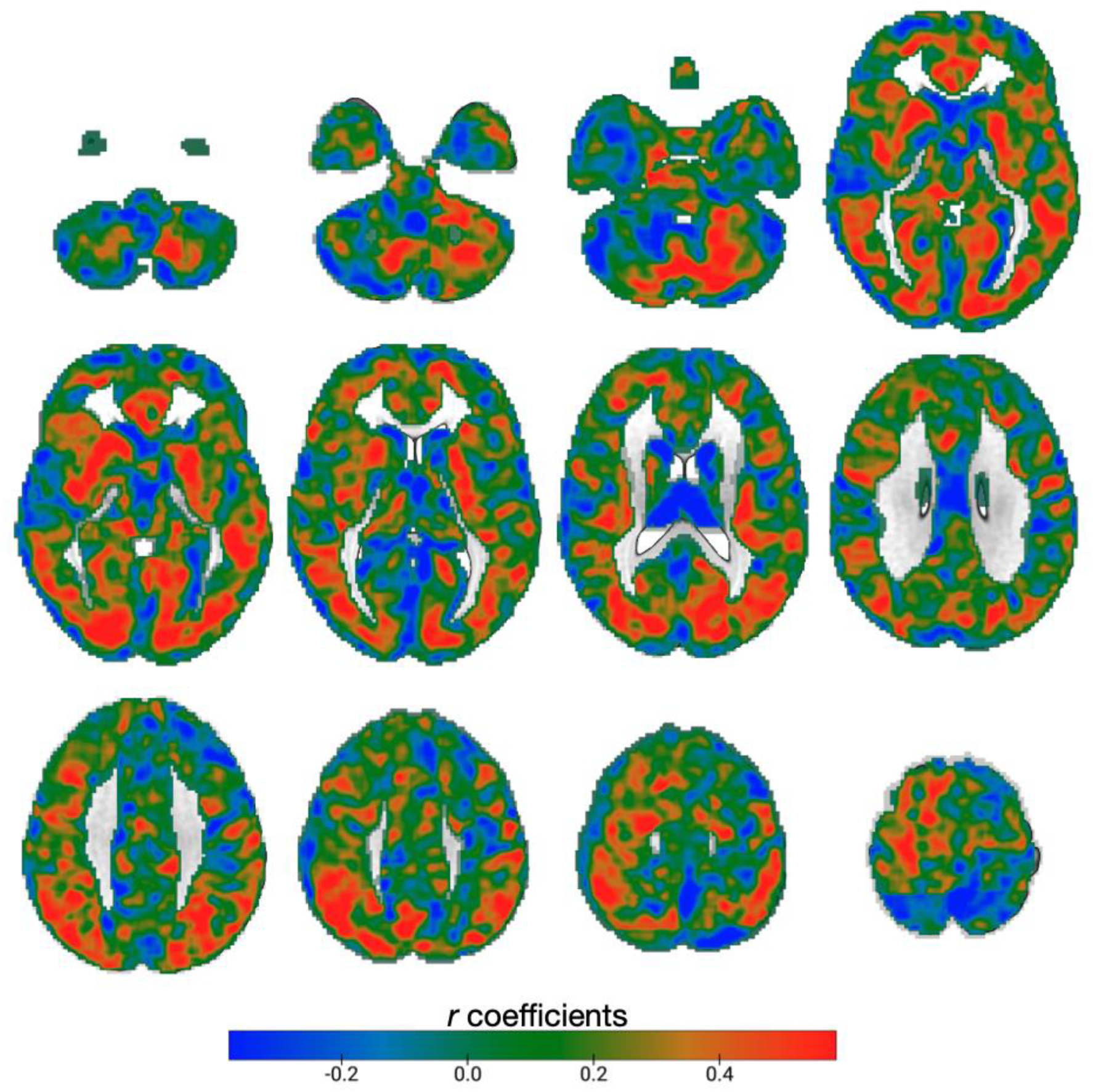
Unthresholded brain-wide correlations between resting state amplitude of low frequency fluctuations and SV2A distribution in control subjects: r-coefficient statistical images representative of brain-wide correlations between amplitude of low frequency fluctuations at rest and [11C]UCB-J DVR_CS_ in healthy control subjects (*n*=16). Voxel-wise Pearson’s correlations, uncorrected. Axial slices shown in MNI152 are: -52 -42 -32-22-12; -2 8 18 28; 38 48 56 66.

**Supplementary figure 2:**
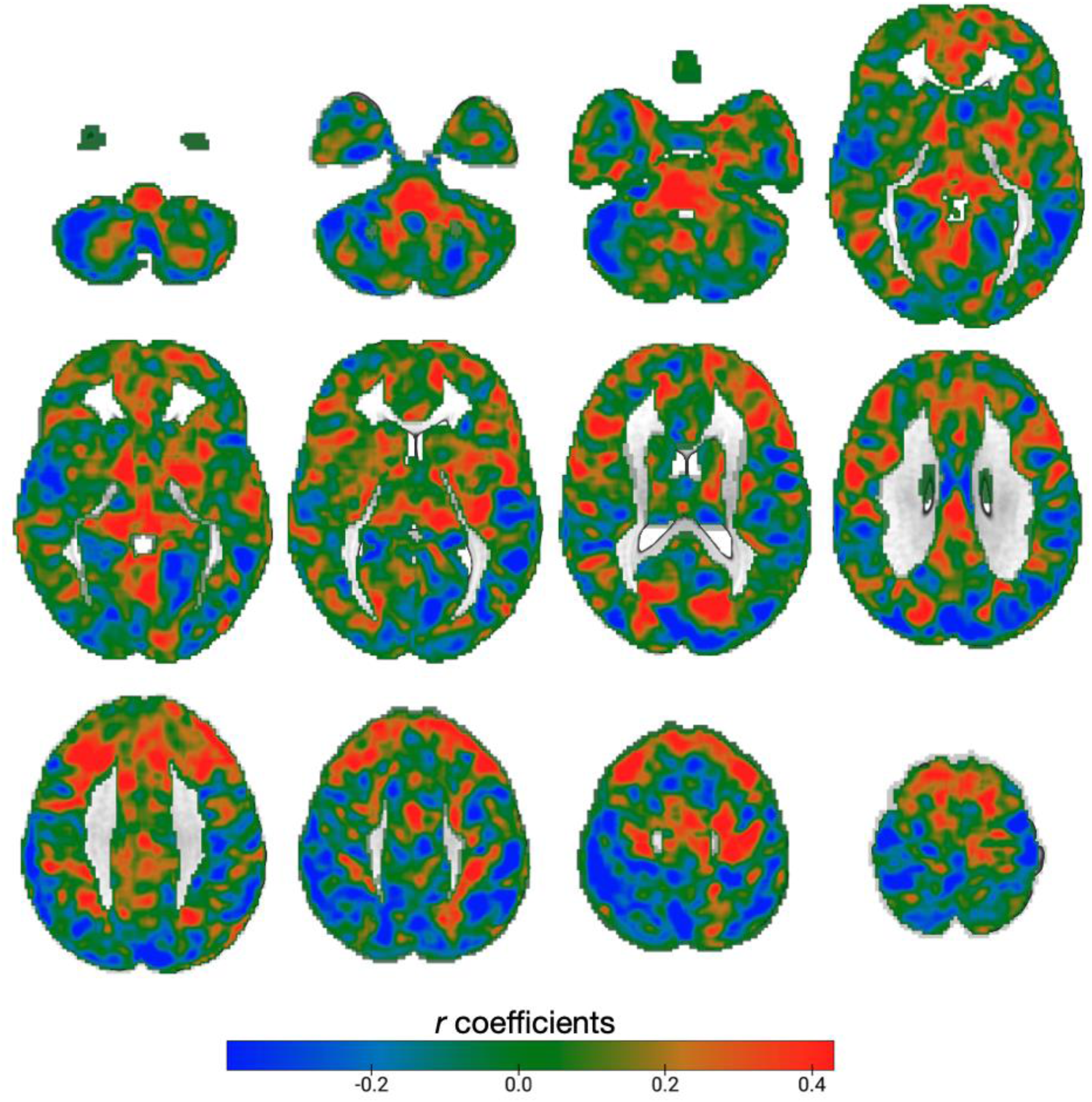
Unthresholded brain-wide correlations between resting state amplitude of low frequency fluctuations and SV2A distribution in patients with schizophrenia: r-coefficient statistical images representative of brain-wide correlations between amplitude of low frequency fluctuations at rest and [11C]UCB-J DVR_CS_ in patients with schizophrenia not taking any medication (*n*=22). Voxel-wise Pearson’s correlations, uncorrected. Axial slices shown in MNI152 are: -52 -42 -32 -22-12; -2 8 18 28; 38 48 56 66.

**Supplementary table 1:**
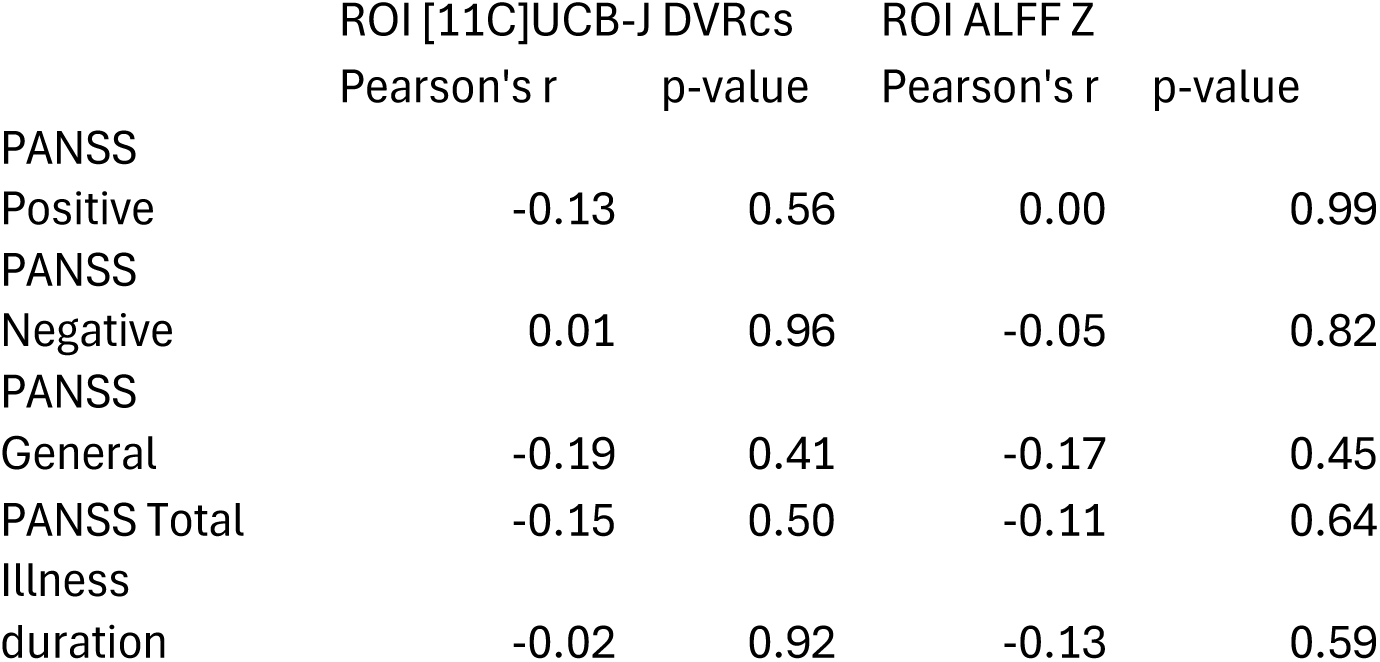

